# Live Cell Extrusion in Cervical Cancer—A Novel Mechanism for Cancer Progression

**DOI:** 10.1101/2025.05.13.653891

**Authors:** Leanna Rose, Sudhir Krishna

## Abstract

Extrusion during development or of transformed cells in normal epithelia has been described as a process of elimination. However, the role of extrusion in transformed epithelia is still unclear. Here, we report that in a primary tumor-derived cervical cancer cell line, SiHa, overcrowding results in the extrusion of a subset of cells. We find that the mechanism of extrusion in SiHa cells is similar to those that drive live cell extrusion in normal epithelia, suggesting the utility of this model. We propose that the subset that is extruded during overcrowding is resistant to anoikis and acquires promigratory features. We find that this population also exhibits an increase in TGF-β signaling, which we show is a promigratory factor in cervical cancer cells and is a potential driver for the migratory potential observed in the extruded population. Our study shows that extrusion in cervical cancers in response to overcrowding underlies promigratory behavior of sub-populations in cervical cancer cell lines.

## Introduction

Cell extrusion is a fundamental biological process wherein cells are expelled from epithelial layers, playing a critical role in tissue homeostasis and integrity. In normal epithelia, this mechanism ensures the removal of apoptotic, damaged, or overcrowded cells without compromising barrier function. The process involves the formation of an actomyosin contractile ring around the cell destined for expulsion, generating the mechanical force necessary to squeeze the cell out of the epithelial layer (Rosenblatt et al., 2001; Slattum & Rosenblatt, 2014). Two primary molecular pathways govern this process: sphingosine-1-phosphate (S1P) signaling through S1P receptor 2 (S1PR2) and mechanotransduction via the Piezo1 ion channel, both of which orchestrate the cytoskeletal rearrangements essential for extrusion (Gu et al., 2011; Gudipaty et al., 2017).

The dual nature of cell extrusion in cancer biology has emerged as an area of significant interest. Initially characterized as a tumor-suppressive mechanism termed Epithelial Defense Against Cancer (EDAC), whereby normal epithelial cells recognize and extrude transformed cells to prevent malignant growth (Kajita et al., 2014), cell extrusion has paradoxically been implicated in facilitating cancer progression. Recent evidence suggests that extrusion may provide an escape route for viable cancer cells from the primary tumor, potentially contributing to early dissemination and metastasis (Slattum & Rosenblatt, 2014). This duality parallels the concept of cell competition, where cells with different fitness levels compete for survival within tissues. In cancer contexts, this competition can result in the selective extrusion of cells with distinct oncogenic properties, influencing tumor heterogeneity and progression (Wagstaff & Kolahgar, 2021).

Cervical cancer, predominantly caused by human papillomavirus (HPV) infection, represents a significant global health burden with high mortality rates in developing countries. The progression from HPV infection to invasive carcinoma involves complex molecular alterations, including the integration of viral DNA and expression of viral oncoproteins E6 and E7, which disrupt key tumor suppressor pathways (Hoppe-Seyler et al., 2018). Despite advances in understanding the molecular pathogenesis of cervical cancer, the mechanisms underlying metastatic dissemination remain incompletely elucidated. Conventional models of metastasis emphasize late-stage events in tumor progression; however, emerging evidence suggests that early dissemination of subpopulations with enhanced survival and migratory capabilities may contribute significantly to metastatic spread (Klein, 2009).

The role of transforming growth factor-beta (TGFβ) signaling in cervical cancer progression has been established, with TGFβ promoting epithelial-to-mesenchymal transition (EMT), invasion, and metastasis. However, the potential intersection between TGFβ signaling, cell extrusion, and metastatic dissemination in cervical cancer has not been thoroughly investigated. Furthermore, while cell extrusion has been studied in various epithelial cancers, its specific contribution to cervical cancer progression remains unexplored.

This study investigates the hypothesis that cell extrusion under overcrowding stress conditions may serve as a mechanism for the dissemination of viable cancer cells with enhanced metastatic potential in cervical cancer. We aim to characterize the extrusion process, identify the key molecular pathways involved, and determine the phenotypic properties of extruded cells that may contribute to metastatic progression. Understanding these mechanisms could reveal novel therapeutic targets to prevent metastatic dissemination in cervical cancer.

## Materials and Methods

### Cell Lines and Culture Conditions

SiHa and CaSki cervical cancer cell lines were obtained from American Type Culture Collection (ATCC, Manassas, VA) and maintained in Dulbecco’s Modified Eagle’s Medium (DMEM; Thermo Fisher Scientific, cat# 11965092) supplemented with 10% fetal bovine serum (FBS; Thermo Fisher Scientific, cat# 16000044). Cells were cultured at 37°C in a humidified atmosphere containing 5% CO₂. All experiments were performed using cells within 25 passages from the original ATCC stock to minimize phenotypic drift. Cultures were tested for mycoplasma contamination biweekly using the MycoAlert Mycoplasma Detection Kit (Lonza, cat# LT07-318) and consistently tested negative throughout the study period. Cell morphology and growth characteristics were routinely monitored by phase-contrast microscopy to ensure the maintenance of expected phenotypic properties.

### Cellular Overcrowding Assay

Single-cell suspensions of SiHa cells were prepared by trypsinization of subconfluent monolayers followed by gentle trituration. Cells were counted using a hemocytometer and seeded in six-well tissue culture plates (Corning, cat# 3516) at a density of 0.5 × 10^6 cells/ml in DMEM supplemented with 10% FBS. Cultures were maintained until they reached confluence (designated as “X” confluence), at which point the medium was replaced with fresh DMEM containing 10% FBS. Cells were then allowed to continue proliferating for approximately 72 additional hours until they reached three times the confluent density (designated as “3X” confluence), as determined by cell counting in representative fields.

For collection of extruded cells, 3X confluent monolayers were washed twice with phosphate-buffered saline (PBS, pH 7.4) to remove non-adherent cells, and fresh medium was added with or without the following compounds: gadolinium chloride (GdCl₃, 10 μM; Sigma-Aldrich, cat# 439770), JTE013 (10 μM; Cayman Chemical, cat# 10009458), SB431542 (10 μM; Selleck Chemicals, cat# S1067), or recombinant human TGFβ1 (5 ng/ml; R&D Systems, cat# 240-B). After 24 hours of incubation, culture supernatants containing extruded cells were collected, and cells were pelleted by centrifugation at 300 × g for 5 minutes. Cell numbers were quantified using a hemocytometer, and viability was assessed by trypan blue exclusion. For microscopic analysis, cultures were observed using a Nikon ECLIPSE TE2000-S inverted microscope equipped with a digital camera (Nikon DS-Qi2) and NIS-Elements software (Nikon).

### Cell Migration Assays

#### Transwell Migration Assay

Transwell migration assays were performed using 8.0 μm pore size polycarbonate membrane inserts (Corning, cat# 3422) in 24-well plates. SiHa cells (5 × 10^4) were suspended in 100 μl serum-free DMEM and placed in the upper chamber, while the lower chamber contained 600 μl DMEM supplemented with 10% FBS as a chemoattractant. After 24 hours of incubation at 37°C, non-migrated cells on the upper surface of the membrane were removed using a cotton swab. Cells that had migrated to the lower surface were fixed with 4% paraformaldehyde for 15 minutes and stained with 0.1% crystal violet for 20 minutes. The membranes were then washed thoroughly with PBS and mounted on glass slides. Migrated cells were counted in ten random fields per membrane using a Nikon ECLIPSE TE2000-S microscope at 200× magnification. Each condition was tested in triplicate, and experiments were repeated at least three times.

#### Wound Healing Assay

SiHa cells were seeded in 6-well plates and grown to 100% confluence. A sterile 200 μl pipette tip was used to create a linear scratch across the cell monolayer. Detached cells were removed by washing twice with PBS, and fresh medium containing 1% FBS (to minimize proliferation) was added. Phase-contrast images of the scratch area were captured immediately (0 hours) and at 24 hours post-scratch using a Nikon ECLIPSE TE2000-S inverted microscope. The scratch width was measured at ten random positions along the wound for each timepoint using ImageJ software (NIH, version 1.53). Wound closure was calculated as the percentage of the initial wound area that was covered by cells after 24 hours. Each experiment was performed in triplicate and repeated at least three times.

#### Anchorage-Independent Growth Assay

Soft agar colony formation assays were performed to assess anchorage-independent growth. Six-well plates were prepared with a base layer of 0.6% agar (Sigma-Aldrich, cat# A1296) in DMEM supplemented with 10% FBS. SiHa cells (5 × 10^3 per well) were suspended in 0.3% agar in DMEM containing 10% FBS and layered over the solidified base layer. The plates were incubated at 37°C in a humidified atmosphere with 5% CO₂ for 21 days, with 500 μl of fresh medium added to the surface of each well twice weekly to prevent desiccation. Colonies (defined as cell clusters with ≥50 cells) were counted in ten randomly selected fields per well under a Nikon ECLIPSE TE2000-S microscope at 100× magnification. Each experimental condition was tested in triplicate, and experiments were repeated at least three times.

#### Statistical Analysis

All quantitative data are presented as mean ± standard error of the mean (SEM) from at least three independent biological replicates (n ≥ 3), unless otherwise specified. Statistical significance was determined using GraphPad Prism software (version 9.0). For comparisons between two groups, two-tailed unpaired Student’s t-tests were applied. For multiple group comparisons, one-way analysis of variance (ANOVA) followed by Tukey’s post hoc test was used. Differences were considered statistically significant at p < 0.05. The specific number of replicates for each experiment is indicated in the corresponding figure legends.

#### Data Availability

The RNA sequencing data generated during this study have been deposited in the NCBI Gene Expression Omnibus (GEO) database under accession number [to be added upon submission]. All other datasets supporting the findings of this study are included within the manuscript and its supplementary information files or are available from the corresponding author upon reasonable request.

## Results

### Overcrowding stress leads to live cell extrusion in primary tumor-derived cervical cancer cells

SiHa cells, derived from a primary cervical cancer lesion, were subjected to overcrowding conditions to investigate cell extrusion behavior. SiHa monolayers grown to 100% confluence (X confluence, approximately 7×10^6 cells per T75 flask) were allowed to continue growing until they reached approximately 3X confluence (approximately 21×10^6 cells per T75 flask), at which point cells began to be extruded into the medium (Fig. 1a). Quantification of the extruded cells revealed that the majority were viable, with live cell extrusion increasing progressively from 6 hours to 24 hours post-overcrowding, followed by a slight decline at 30 hours (Fig. 1a). The 24-hour time point, which showed maximum extrusion, was used for subsequent experiments.

**Figure 1.**
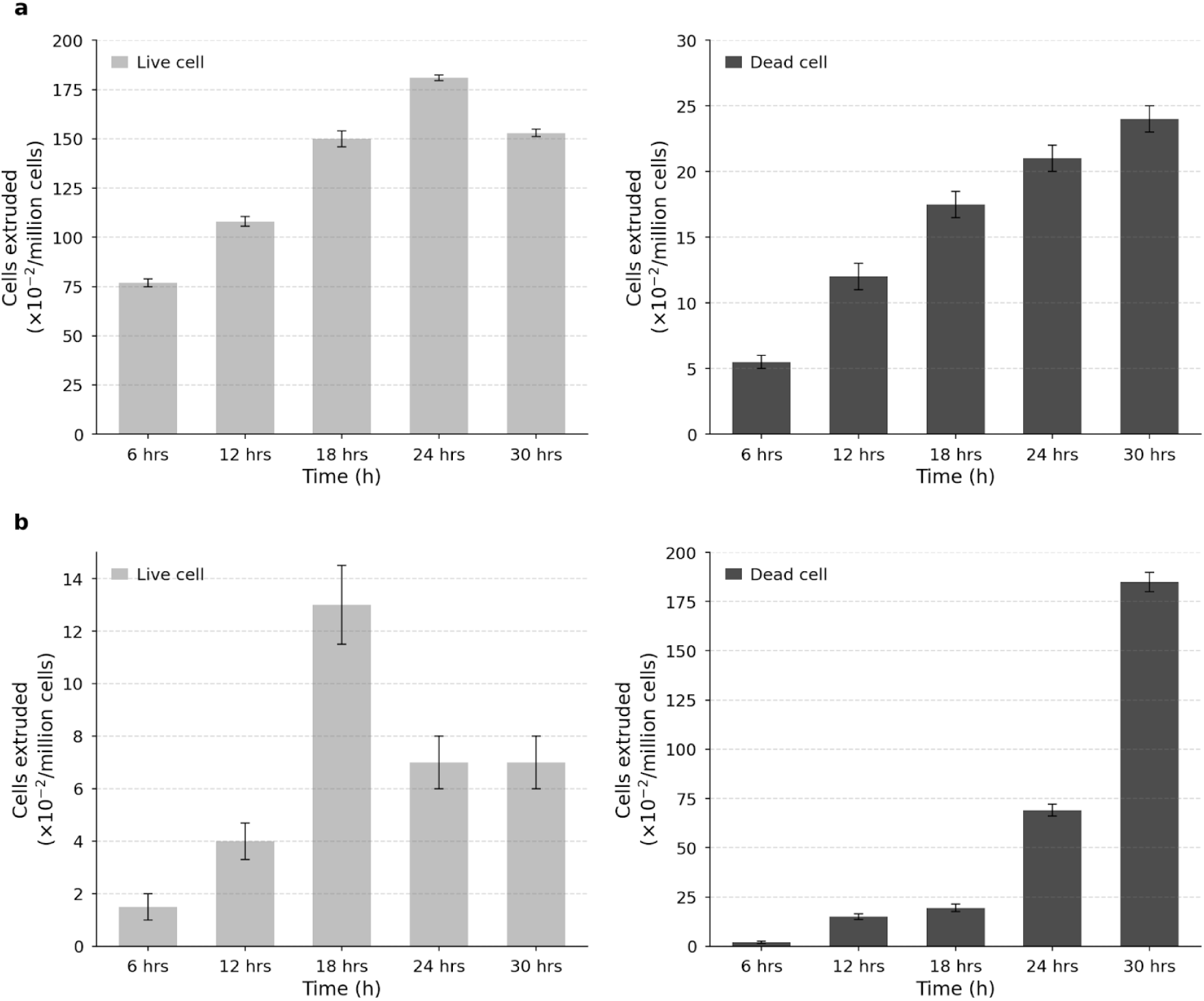
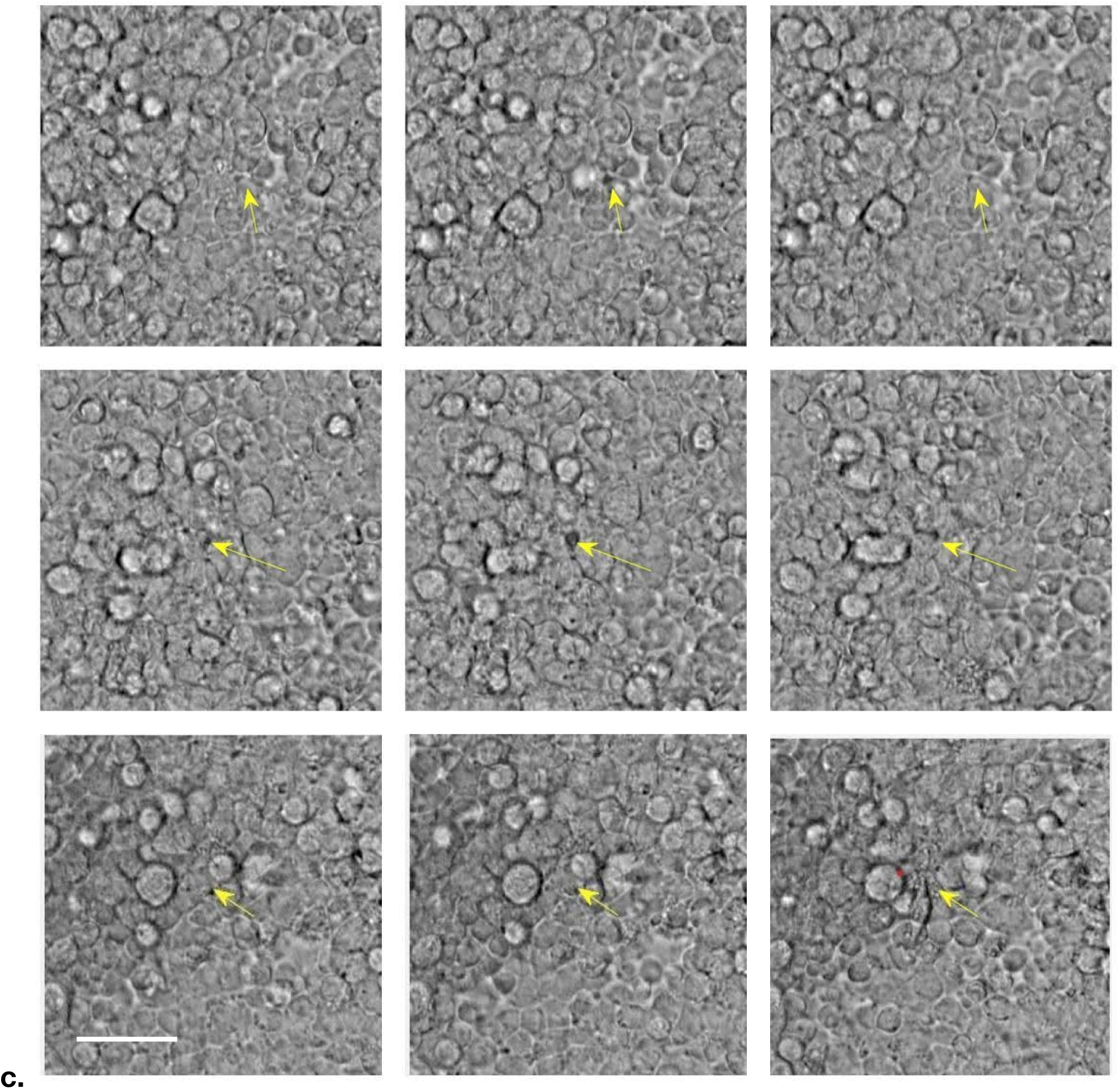
Cell extrusion dynamics in primary tumour-derived cervical cancer cell lines. a, Quantification of extruded cells in SiHa cultures at different time points. Live cell extrusion (left, light gray) increases over time, peaking at 24 h, while dead cell extrusion (right, dark gray) shows a steady increase. b, Quantification of extruded cells in CasKi cultures demonstrates different extrusion dynamics compared to SiHa cells. Live cell numbers(left) remains relatively low while dead cell numbers (right) increases dramatically at later time points, suggesting different mechanisms of extrusion between the two cell lines. c, Time-lapse phase-contrast microscopy tracking the extrusion process in SiHa cells. Yellow arrows indicate a cell undergoing extrusion from the monolayer over a 20-minute period. All quantification data represent mean ± s.e.m. from n = 3 biologically independent experiments. Statistical analysis was performed using one-way ANOVA with Tukey’s multiple comparisons test. Scale bar, 20 μm.

In contrast, CaSki cells, derived from a metastatic cervical cancer lesion, displayed markedly different behavior under overcrowding stress. Unlike SiHa cells, CaSki cultures showed a visibly disrupted monolayer with cell loss in patches, and the extruded cells were predominantly non-viable (Fig. 1b). This distinct extrusion pattern between primary tumor-derived and metastasis-derived cell lines suggests fundamental differences in how these cells respond to tissue density stress.

Time-lapse microscopy of overcrowded SiHa monolayers revealed both apical and basal extrusion events (Fig. 1c). Notably, we observed instances where apically extruded cells subsequently displayed active migration on top of the monolayer, suggesting retention of migratory capability following extrusion.

### Live cell extrusion in SiHa cells is regulated by S1P signaling and Piezo1 activation

To validate the molecular mechanisms underlying live cell extrusion in SiHa cells, we investigated key pathways previously implicated in epithelial cell extrusion. Treatment with Gdף+, an inhibitor of stretch-activated channels including Piezo1, significantly reduced live cell extrusion in a dose-dependent manner (Fig. 2a). Notably, Gdף3+ treatment resulted in the formation of cell masses on the monolayer, suggesting impaired extrusion processes.

**Figure 2.**
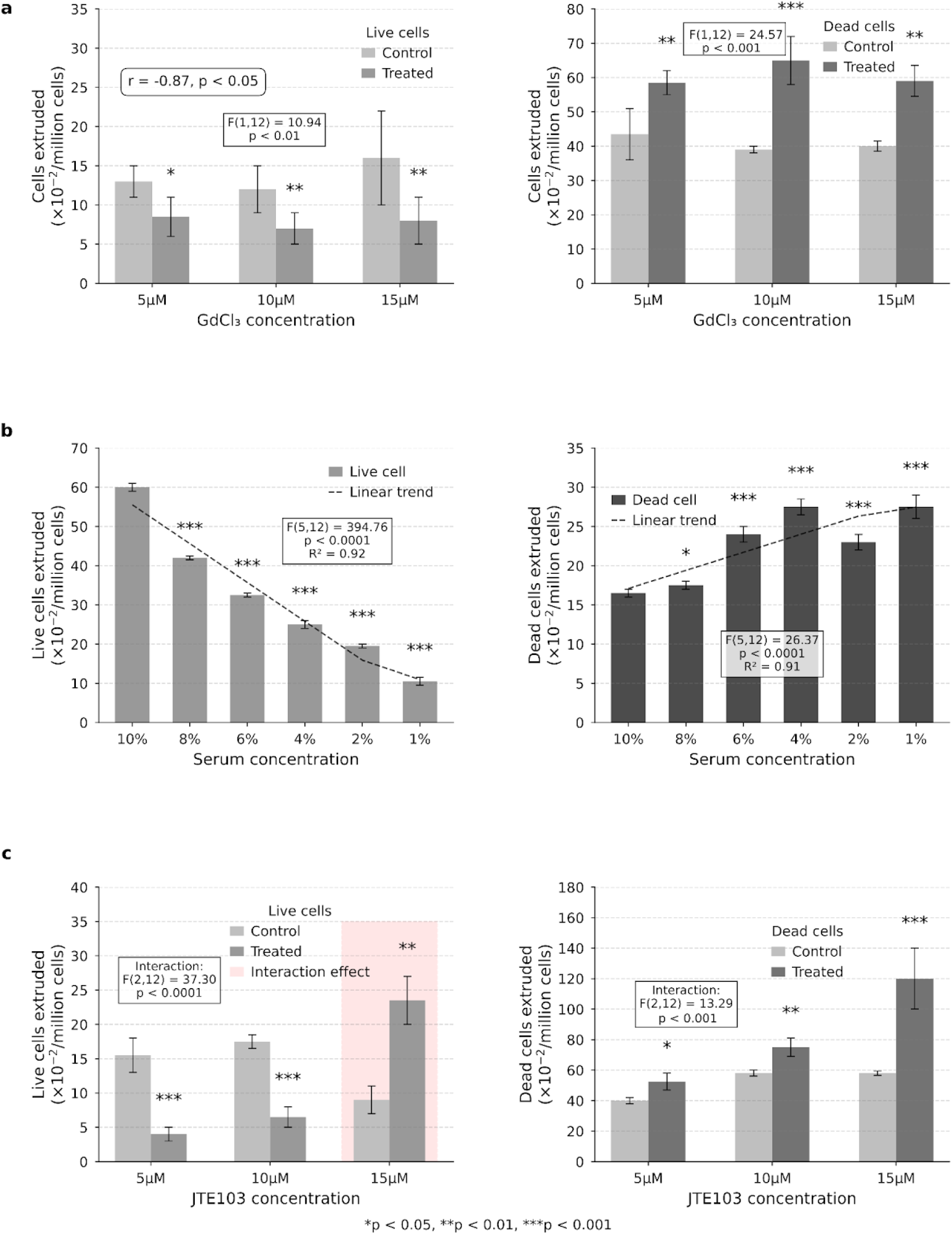
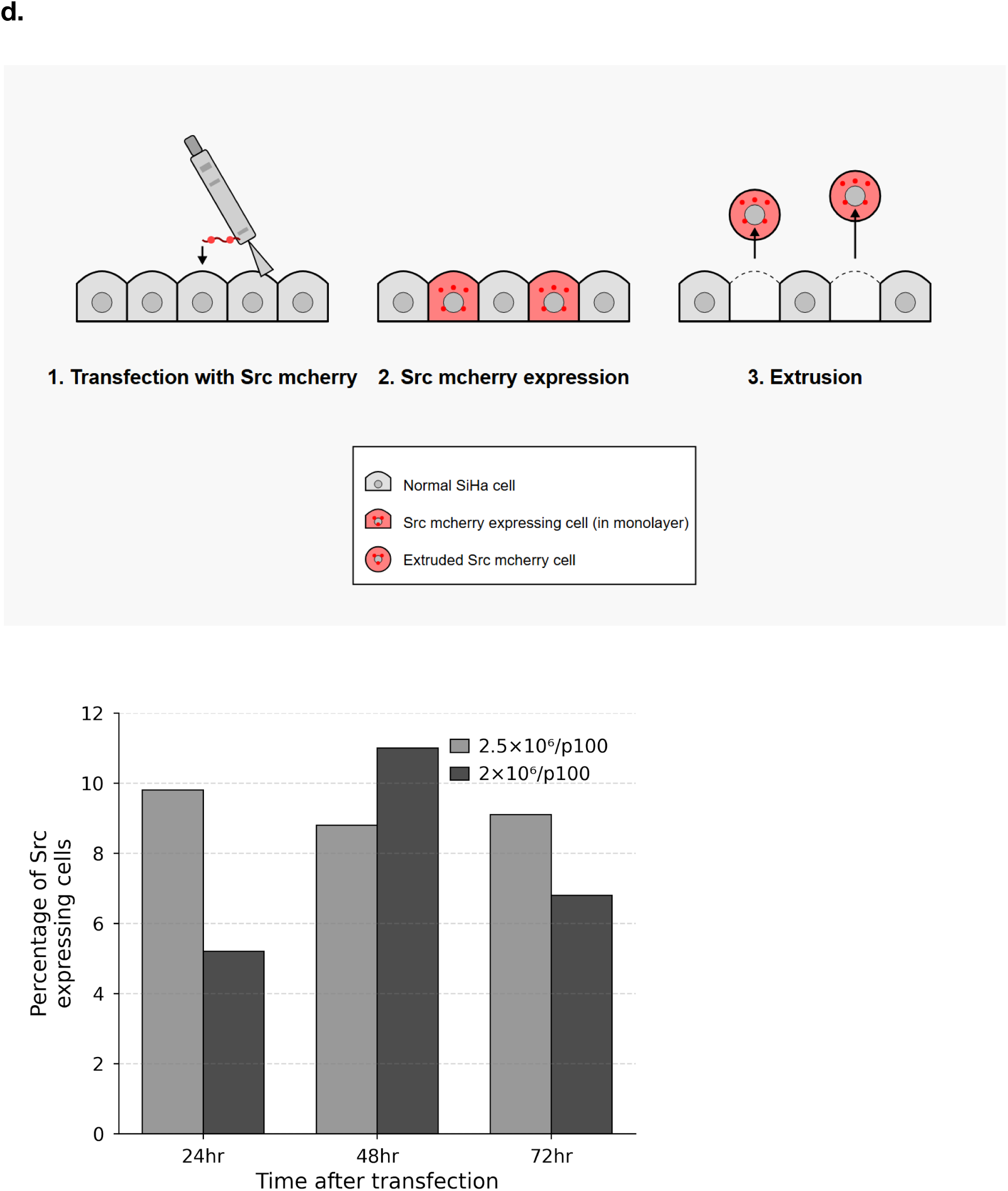
Cell extrusion is regulated by mechanosensitive channels and sphingolipid signaling. a, Effects of the Piezo1 inhibitor GdCl₃ on cell extrusion. Treatment with GdCl₃ (5-15μM) significantly reduced live cell extrusion (F(1,12) = 10.94, p < 0.01, left) while increasing dead cell extrusion (F(1,12) = 24.57, p < 0.001, right). A strong negative correlation was observed between live and dead cell extrusion rates (r = -0.87, p < 0.05). b, Serum withdrawal progressively alters cell extrusion patterns. Decreasing serum concentration from 10% to 1% showed a significant linear trend in reducing live cell extrusion (F(5,12) = 394.76, p < 0.0001, R² = 0.92, left) while enhancing dead cell extrusion (F(5,12) = 26.37, p < 0.0001, R² = 0.91, right). All reduced serum conditions differed significantly from the 10% control (p < 0.001). c, Effects of S1PR2 inhibitor JTE103 on cell extrusion. JTE103 treatment showed a significant interaction effect between concentration and treatment for both live (F(2,12) = 37.30, p < 0.0001) and dead cells (F(2,12) = 13.29, p < 0.001). For live cells, treatment reduced extrusion at lower concentrations (5-10μM, p < 0.001) but significantly increased it at 15μM (p < 0.01). Dead cell extrusion was consistently enhanced across all concentrations tested, with the most pronounced effect at 15μM (p < 0.001). d, Percentage of Src-mCherry expressing cells in SiHa monolayers over time. Cells were seeded at two different densities (2.5×10⁶/p100 and 2×10⁶/p100) and monitored at 24, 48, and 72 hours after transfection. No significant changes were observed over time, suggesting a dynamic equilibrium between Src expression and cell extrusion. Data represent mean ± SEM from n=3 independent experiments.

Since sphingosine-1-phosphate (S1P) signaling has been reported to mediate cell extrusion, we examined its role in our system through two approaches. First, serum withdrawal experiments, which reduce bioavailable S1P, showed a progressive decrease in live cell extrusion with decreasing serum concentrations (Fig. 2b). Second, treatment with JTE013, a specific inhibitor of S1P receptor 2 (S1PR2), similarly reduced live cell extrusion in a dose-dependent manner (Fig. 2c). These results confirm that S1P signaling through S1PR2 is critical for live cell extrusion in SiHa cells.

Previous studies in MDCK cells have shown that transformed cells overexpressing oncogenes like Ras or Src are selectively extruded from epithelial monolayers. To determine if SiHa monolayers exhibit similar selective extrusion, we transfected a small percentage of cells with Src-mCherry and monitored their fate. Analysis of the extruded cell population at 24, 48, and 72 hours revealed progressive enrichment of Src-positive cells (Fig. 2d), indicating that SiHa monolayers preferentially extrude cells with higher Src expression.

### Extruded SiHa cells exhibit enhanced anoikis resistance and migratory potential

To characterize the phenotypic properties of extruded cells, we collected and replated cells extruded from overcrowded SiHa monolayers. A subset of these extruded cells successfully re-adhered and proliferated, which we designated as RE1 cells. As a control, we similarly subcultured cells from the monolayer (RM1 cells) (Fig. 3a).

**Figure 3.**
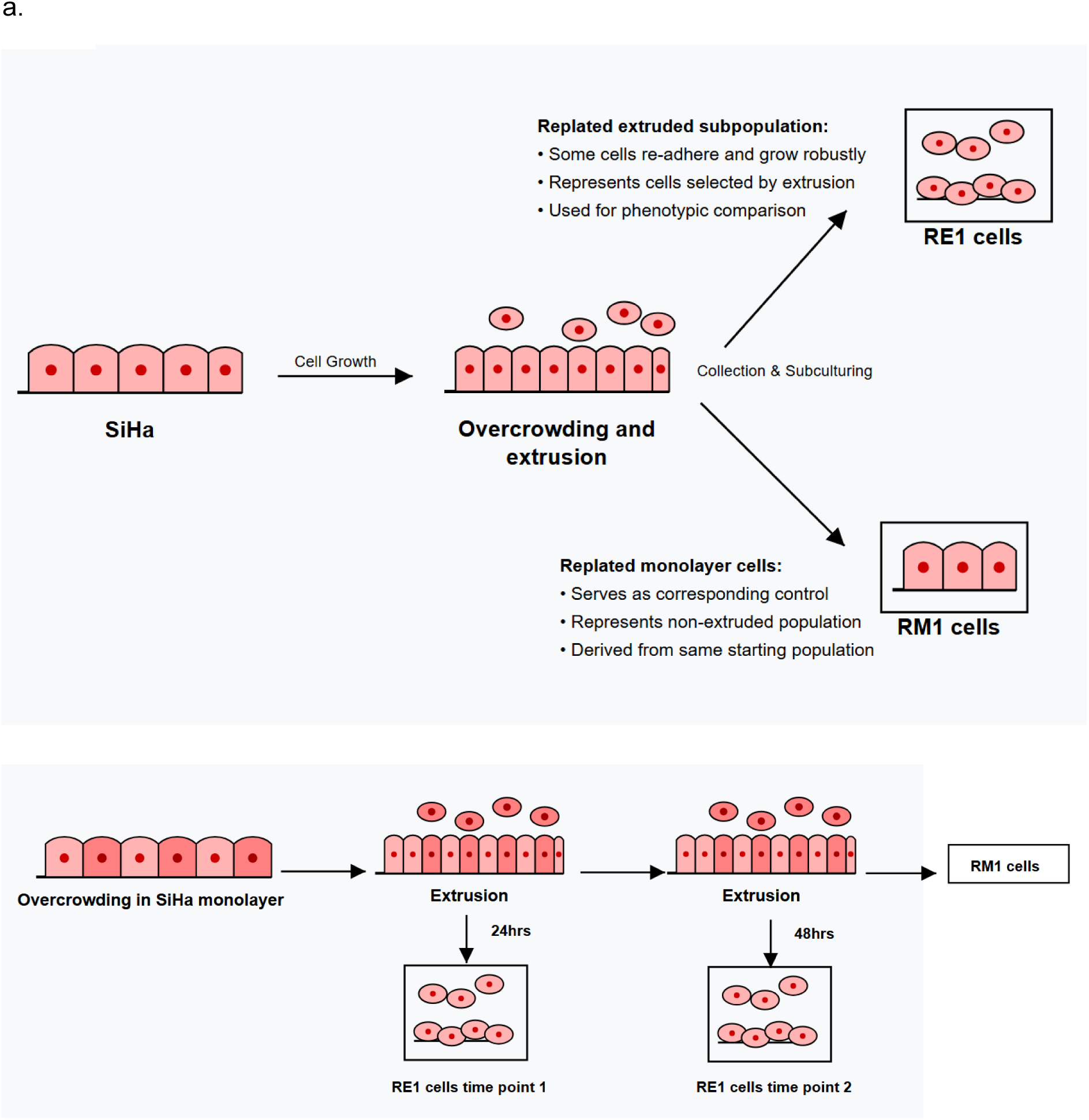
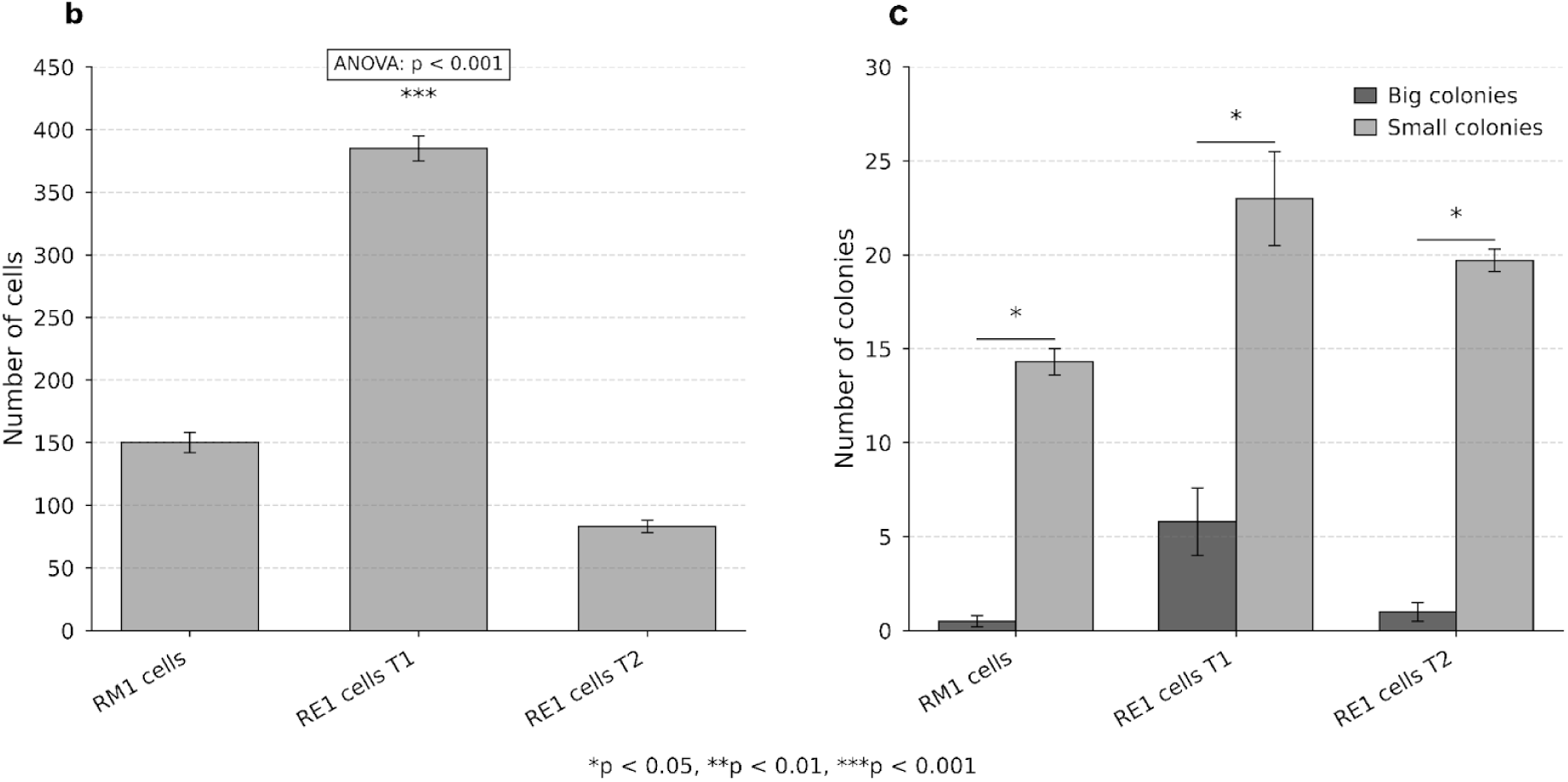
Phenotypic characterization of the extruded subset in SiHa. Cellular characteristics and colony-forming ability of RM1 and RE1 cell lines. a,Schematic showing the subculturing of extruded subset. b, Cell number comparison between RM1 cells and two RE1 cell subpopulations (T1 and T2) in a transwell migration assay. RE1 cells T1 show significantly higher cell counts (385 ± 10) compared to both RM1 cells (150 ± 8) and RE1 cells T2 (83 ± 5) (one-way ANOVA, p < 0.001). c, Colony formation assay demonstrating differential distribution of big and small colonies across cell types. RE1 cells T1 exhibit the highest colony-forming efficiency for both big colonies (5.8 ± 1.8) and small colonies (23.0 ± 2.5), with significant differences between colony sizes (p < 0.01). RM1 cells and RE1 cells T2 show similar patterns with predominant formation of small colonies (14.3 ± 0.7 and 19.7 ± 0.6, respectively) and minimal development of big colonies. Data represent mean ± SEM from three independent experiments.

Soft agar colony formation assays were performed to assess anoikis resistance. RE1 cells formed significantly more colonies than RM1 cells, with particular enrichment in medium-sized colonies (Fig. 3c). This enhanced anchorage-independent growth capability suggests that extruded cells possess greater resistance to anoikis.

The migratory potential of these cells was evaluated using transwell migration assays. RE1 cells displayed markedly enhanced migration compared to RM1 cells, with approximately 2.5-fold more cells traversing the membrane in response to a chemotactic gradient (Fig. 3b). This increased migratory capability suggests that extruded cells may have greater potential for invasion and metastasis.

To distinguish whether these phenotypic differences arose from pre-existing heterogeneity or were induced by the extrusion process, we collected and characterized cells extruded at different time points: 24 hours (RE1) and 48 hours (designated as RE1 time point 2). While both RE1 and RE1 time point 2 cells showed comparable anoikis resistance in soft agar assays, their migratory capabilities differed considerably. RE1 cells showed substantially higher migration than both RM1 and RE1 time point 2 cells. This temporal difference suggests that enhanced anoikis resistance may be induced by the extrusion process itself, while the migratory phenotype might reflect pre-existing heterogeneity within the monolayer, with more migratory cells being preferentially extruded earlier.

### TGFβ signaling drives the migratory phenotype in extruded cells

To investigate the molecular basis of the enhanced migratory phenotype in extruded cells, we performed RNA sequencing analysis comparing freshly extruded cells to monolayer cells which revealed a bulk downregulation of mRNA transcripts in the extruded population, with selective upregulation of genes associated with oxidative phosphorylation (Supplementary Fig. 1a). This expression pattern resembles cellular stress responses observed in various contexts.

To further investigate these metabolic alterations, we performed mitotracker green staining and observed increased mitochondrial mass in RE1 cells compared to RM1 cells (Supplementary Fig. 1b). However, Seahorse assays revealed reduced oxygen consumption rates in RE1 cells (Supplementary Fig. 1c), suggesting a shift toward glycolytic metabolism despite increased mitochondrial content.

To better understand the molecular pathways involved in the migratory phenotype observed in extruded cells, we performed RNA sequencing analysis comparing RE1 cells to RM1cells which revealed an upregulation of TGFβ signalling (Supplementary Fig. 2). We then examined the role of TGFβ signalling, a known driver of cell migration. Treatment of SiHa cells with SB431542, a selective inhibitor of TGFβ receptor type I, significantly reduced live cell extrusion in a dose-dependent manner (Fig. 4a). This suggests that TGFβ signalling contributes to the extrusion process in SiHa cells.

**Figure 4.**
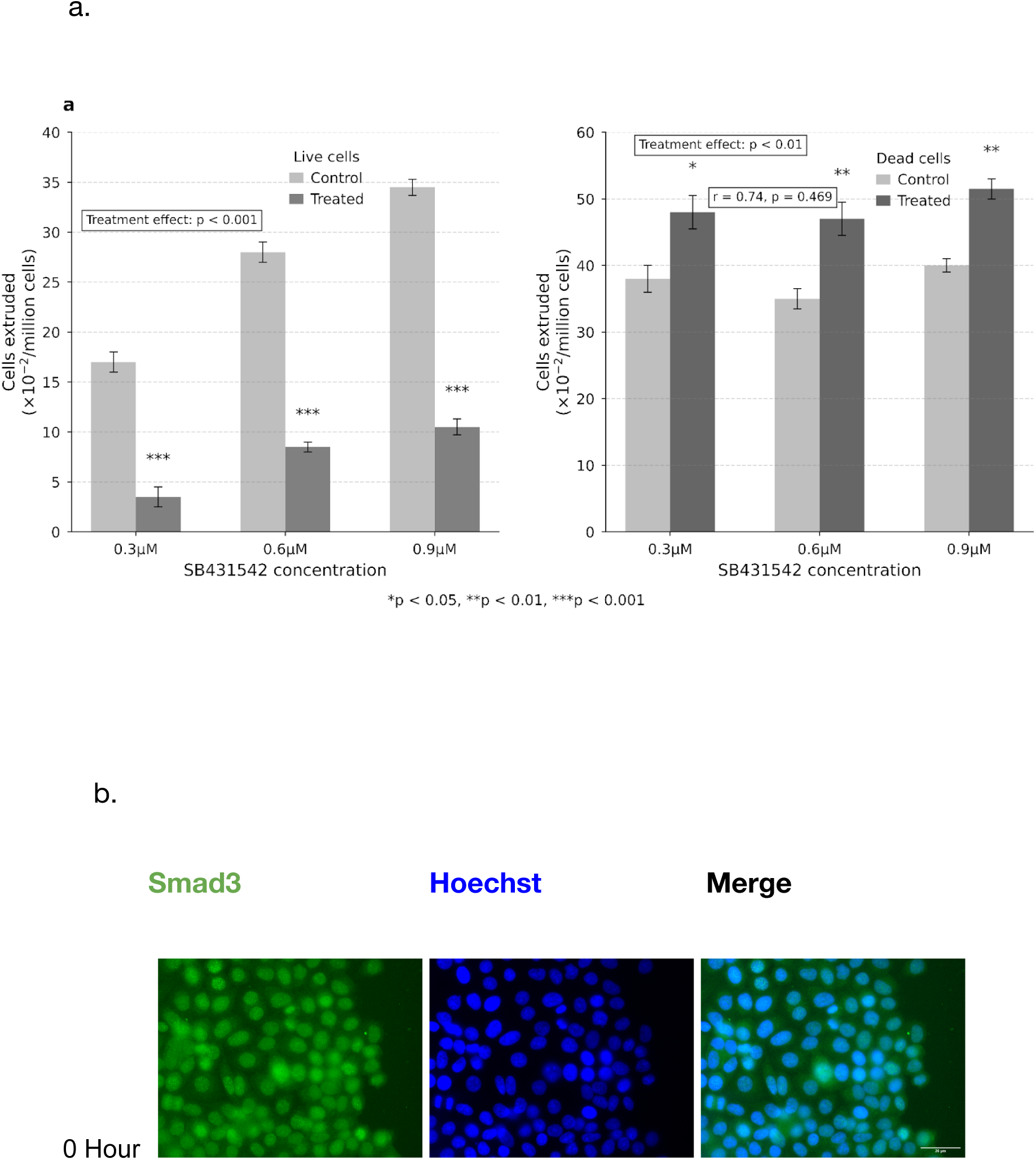
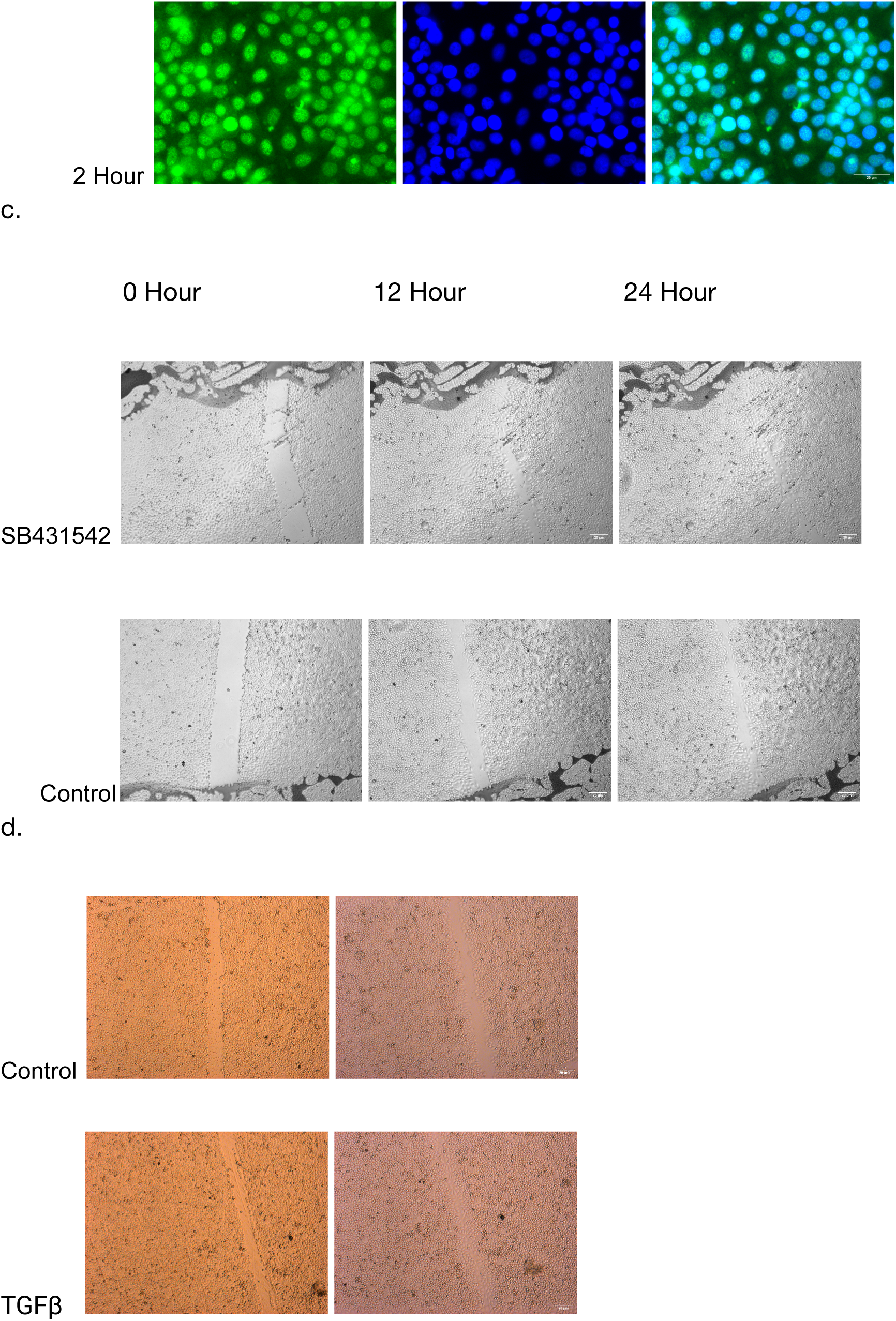
TGFβ signalling a driver for the migratory phenotype observed in RE1 cells. a, SB431542 (TGFβ inhibitor) differentially affects live and dead cell extrusion in a dose-dependent manner. Live cell extrusion exhibits a concentration-dependent response, with the inhibitory effect increasing from 0.3μM (3.5 ± 1.0 cells) to 0.9μM (10.5 ± 0.8 cells). Treatment effect is strongest at the lowest concentration, with a 79% reduction in live cell extrusion compared to control. b, In contrast, SB431542 treatment increases dead cell extrusion across all concentrations. The effect is statistically significant at each concentration (0.3μM: p < 0.05; 0.6μM: p < 0.01; 0.9μM: p < 0.01), with a slight positive correlation between concentration and extrusion rate. The highest extrusion rate was observed at 0.9μM (51.5 ± 1.5 cells). b, Subcellular localization of Smad3 protein visualized by immunofluorescence microscopy. Left panels show Smad3 immunostaining (green), middle panels show nuclear DNA stained with Hoechst (blue), and right panels show merged images. The top row displays cells at 0 hour (baseline/ohour after scratch wound) where Smad3 shows predominantly cytoplasmic distribution. The bottom row shows cells at a later time point following migration cue stimulation, demonstrating increased nuclear accumulation of Smad3, indicative of active TGF-β signaling and Smad3 nuclear translocation. Scale bar = 50 μm. c,Time-dependent effects of TGF-β signaling inhibition on cell migration in a wound healing assay. Phase-contrast microscopy images of cell monolayers treated with the TGF-β type I receptor inhibitor SB431542 (top row) or vehicle control (bottom row) at 0, 12, and 24 hours after wound creation. Control cells demonstrate progressive wound closure over the 24-hour period, whereas SB431542-treated cells show impaired migration and reduced wound healing capacity, indicating the requirement of TGF-β signaling for effective cell migration. Images were captured at 10× magnification. d, Effect of TGF-β treatment on cell migration. Bright-field microscopy images comparing untreated control cells (top row) with TGF-β-treated cells (bottom row) in a scratch wound assay. Left and right panels represent different time points 0 hour and 4 hour after scratch wound was created. Control cells demonstrate progressive wound closure over the 4-hour period, whereas TGF-β-treated cells show accelerated migration, indicating the role of TGF-β in regulating epithelial cell motility. Scale bar in the upper right image represents 100 μm.

Immunofluorescence analysis revealed translocation of Smad3, a key mediator of TGFβ signaling, from the cytoplasm to the nucleus in migrating SiHa cells (Fig. 4b). This nuclear localization of Smad3 indicates active TGFβ signaling in these cells.

Scratch wound assays further confirmed the role of TGFβ signaling in SiHa cell migration. Treatment with SB431542 impaired wound closure compared to control conditions (Fig. 4c), while exogenous TGFβ treatment enhanced wound closure (Fig. 4d). These findings demonstrate that TGFβ signaling is a critical driver of the migratory phenotype observed in extruded SiHa cells.

### Extruded cells display altered cellular respiration and transcriptional profiles

To investigate the effect of TGFβ signalling, a known EMT factor on the EMT status of the extruded cells we computed the EMT score for each sample using transcriptome data from RNA sequencing performed on RE1 and RM1 cells and RE1 and RM1 cells primed for migration using a cytokine gradient for 6 hours, to trigger transcriptional changes associated with cell migration, to estimate the EMT phenotype (Tan et al., 2014) (Supplementary Table 2). The EMT scores of both RE1 and Control RM1 cells with and without migration cue were distributed between 0.13 to 0.25 indicating that the EMT status did not differ between the samples (Supplementary Table 2), and both cell populations are EM hybrid in nature with a mesenchymal shift in both RE1 and RM1 cells in response to the migration cue.

We then went on to analyse the EMT status in extruded cells using the EMTome database (Vasaikar et al., 2021). We generated Hazard scores for our cancer type and selected the signatures with statistically significant (pvalue <0.05) hazard ratios. Our samples showed an average of 81% genes expressed in the 22 signatures we examined. With an average of 15% upregulation of genes in the RE1 cells compared to RM1 control and a 17% upregulation in the RE1 primed for migration set compared to RM1 control cells also treated with migration cue (Supplementary Table 3). This suggests that cells undergoing extrusion are experiencing EMT activation, which is a critical process in cancer progression that enables cells to gain migratory and invasive properties.

Further, several signatures showing strong upregulation (e.g., Aiello_et_al_2018 at 40%, Tso_et_al_2015 at 33.33%) are associated with aggressive cancer phenotypes. This suggests that extruded cells may represent a subset with higher metastatic potential, supporting the hypothesis that extrusion contributes to tumor dissemination.

## Discussion

Our study demonstrates that overcrowding stress in primary tumor-derived cervical cancer cells triggers substantial live cell extrusion, a process that may have important implications for cancer progression and metastasis. The stark contrast between SiHa (primary tumor-derived) and CaSki (metastasis-derived) cell lines in their extrusion behavior suggests fundamental differences in how cells at different stages of cancer progression respond to density stress.

### Live cell extrusion as a potential mechanism for cancer cell dissemination

The predominance of live cell extrusion in SiHa cells under overcrowding conditions represents a significant finding, as it suggests that primary cervical tumors may utilize this mechanism to disseminate viable cancer cells into the surrounding microenvironment. This process differs from conventional cell death-triggered extrusion observed in normal epithelia, which primarily serves a homeostatic function. Live cell extrusion could potentially represent an early step in the metastatic cascade, wherein overcrowding in the primary tumor induces the expulsion of viable cells that retain proliferative and migratory capabilities.

Our observation that extruded cells can actively migrate on the monolayer surface following apical extrusion further supports this hypothesis. This behavior could facilitate local invasion and eventual distant metastasis.

The reduced incidence of live cell extrusion in CaSki cells, derived from metastatic lesions, suggests that such cells may have evolved alternative mechanisms for tissue invasion that do not rely on extrusion, or that extrusion is primarily a feature of early-stage disease.

### Molecular mechanisms governing live cell extrusion in cervical cancer

Our findings confirm that established mechanistic pathways of cell extrusion are operational in cervical cancer cells. Specifically, the S1P-S1PR2 signaling axis and Piezo1-mediated mechanotransduction appear to be critical for live cell extrusion in SiHa cells. These pathways have been previously implicated in epithelial cell extrusion in various contexts, but their role in cervical cancer was previously unknown.

The inhibition of extrusion by Gd^3+ treatment indicates that stretch-activated channels, likely including Piezo1, are essential mediators of this process. Piezo1 has been shown to sense mechanical strains in overcrowded epithelia, triggering calcium influx and subsequent cytoskeletal rearrangements necessary for extrusion. Similarly, the dependence on S1P signaling, as demonstrated by the effects of serum withdrawal and JTE013 treatment, aligns with established models where S1P produced by extruding cells activates S1PR2 on neighboring cells to trigger contractile ring formation.

The preferential extrusion of Src-overexpressing cells from SiHa monolayers parallels observations in MDCK cells and suggests that cell competition mechanisms may be at play. This selective extrusion could serve as a quality control mechanism to eliminate potentially harmful transformed cells, but paradoxically may also contribute to the dissemination of more aggressive cell variants.

### Phenotypic alterations in extruded cells with implications for metastasis

Perhaps the most striking aspect of our findings is the phenotypic characterization of extruded SiHa cells. These cells exhibit enhanced anoikis resistance and migratory capacity compared to their monolayer counterparts, two traits critically associated with metastatic potential. The ability to survive in an anchorage-independent manner and migrate toward chemoattractants would significantly advantage these cells during metastatic spread.

Our temporal analysis of extrusion revealed an important distinction: while enhanced anoikis resistance was consistently observed in cells extruded at different time points, migratory capacity was highest in cells extruded earlier. This suggests that anoikis resistance may be induced by the extrusion process itself, potentially through activation of survival pathways during detachment. In contrast, enhanced migratory potential likely reflects pre-existing heterogeneity within the monolayer, with more motile cells being preferentially extruded in earlier phases.

The RNA sequencing data showing bulk downregulation of transcripts with selective upregulation of oxidative phosphorylation genes in extruded cells resembles stress response patterns. Cancer cells often undergo metabolic shifts to adapt to changing environmental conditions, and our findings indicate that extruded cells may adopt a Warburg-like metabolism to support their survival and migratory capabilities.

### TGFβ signaling as a driver of the migratory phenotype

Our investigation into the molecular drivers of the enhanced migratory phenotype in extruded cells identified TGFβ signaling as a critical pathway. TGFβ is well-established as an inducer of EMT and cell migration in various cancer types. The nuclear translocation of Smad3 in migrating cells and the effects of TGFβ pathway modulation on wound closure confirm its active role in SiHa cell migration.

The inhibitory effect of SB431542 on live cell extrusion further suggests that TGFβ signaling may be integral to the extrusion process itself, not merely a driver of post-extrusion phenotypes.

This finding adds to our understanding of the molecular mechanisms governing cell extrusion in cancer contexts and identifies potential therapeutic targets.

The enrichment of EMT gene signatures in extruded cells, particularly under migratory conditions, provides further evidence that these cells may be primed for metastatic behavior. EMT is widely recognized as a critical process in cancer progression, enabling epithelial cancer cells to acquire mesenchymal traits that facilitate invasion and metastasis.

### Clinical implications and future directions

Our findings have several potential clinical implications. First, they suggest that targeting cell extrusion mechanisms, such as the S1P-S1PR2 axis or TGFβ signaling, might represent a novel therapeutic approach to prevent the dissemination of viable cancer cells from primary cervical tumors. Second, these signalling pathways could serve as biomarkers for metastatic potential in cervical cancer.

Future research should focus on validating these findings in primary human cervical cancer specimens and investigating additional molecular pathways that might be involved in extrusion-mediated cancer cell dissemination. The role of the tumor microenvironment in modulating cell extrusion and the fate of extruded cells in vivo also warrant further investigation.

In conclusion, our study reveals that live cell extrusion in response to overcrowding stress may represent an important mechanism for cancer cell dissemination in cervical cancer. The extruded cells exhibit enhanced survival and migratory capabilities that could contribute to metastatic spread (Fig. 5). Understanding and potentially targeting this process could open new avenues for preventing cervical cancer progression.

**Figure 5.**
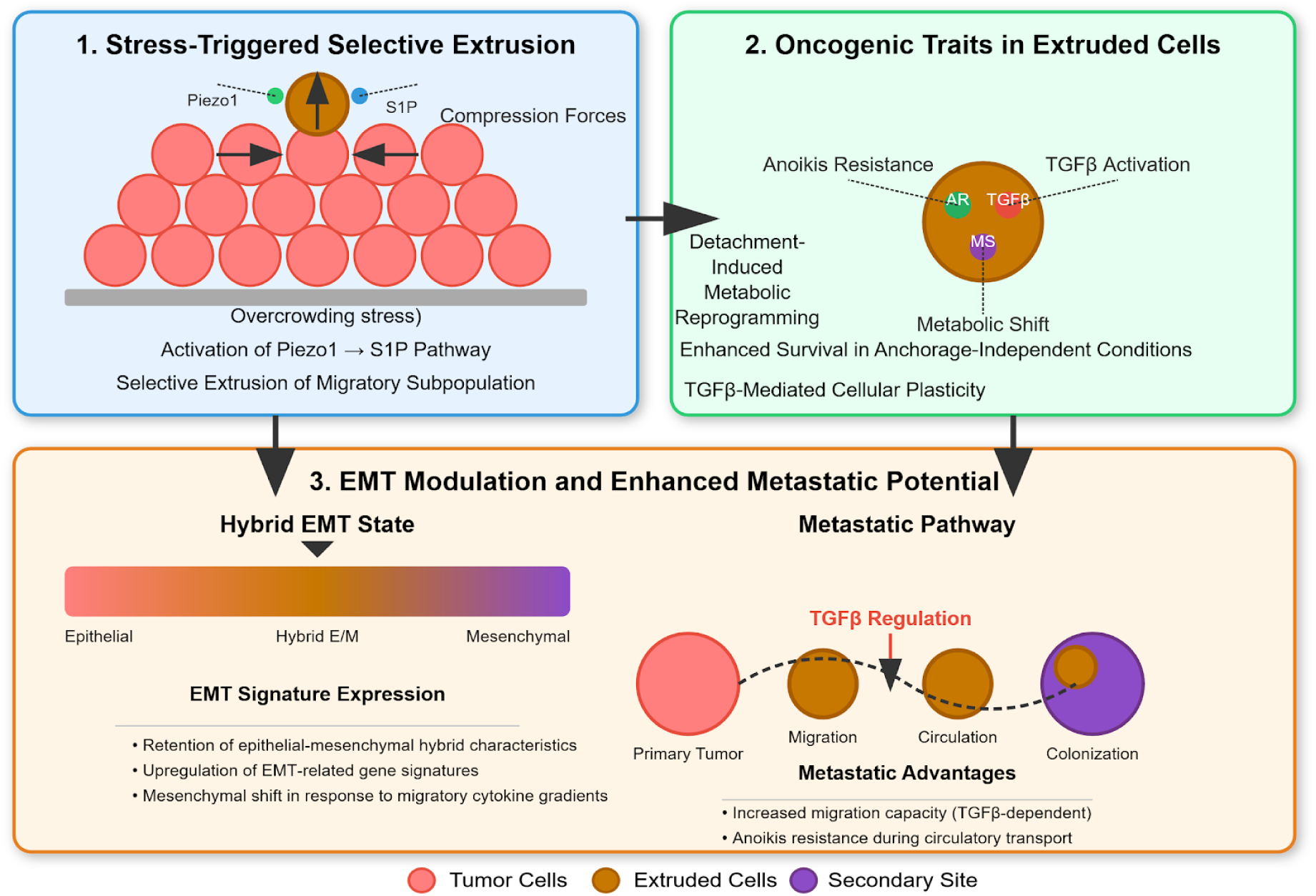
Mechanistic Model of Stress-Triggered Tumor Cell Extrusion and Enhanced Metastatic Potential. This schematic illustrates the proposed three-stage process of tumor cell extrusion and subsequent metastatic progression: Panel 1: Stress-Triggered Selective Extrusion Depicts how mechanical stress from overcrowding activates the Piezo1→S1P (sphingosine-1-phosphate) signaling pathway in tumor cells, leading to selective extrusion of a migratory subpopulation from the primary tumor mass. Panel 2: Oncogenic Traits in Extruded Cells Shows key acquired characteristics of extruded cells (brown), including anoikis resistance, TGFβ activation, metabolic shift (MS), and adaptive reprogramming (AR) that enhance survival in anchorage-independent conditions through TGFβ-mediated cellular plasticity. Panel 3: EMT Modulation and Enhanced Metastatic Potential Illustrates how extruded cells adopt a hybrid epithelial-mesenchymal transition (EMT) state that confers metastatic advantages during migration, circulation, and colonization at secondary sites (purple). The process is regulated by TGFβ signaling throughout the metastatic cascade, promoting migration capacity and survival during circulation. Color code: Primary tumor cells (pink), extruded cells (brown), secondary metastatic site (purple).

---

**Supplementary Figure 1.**
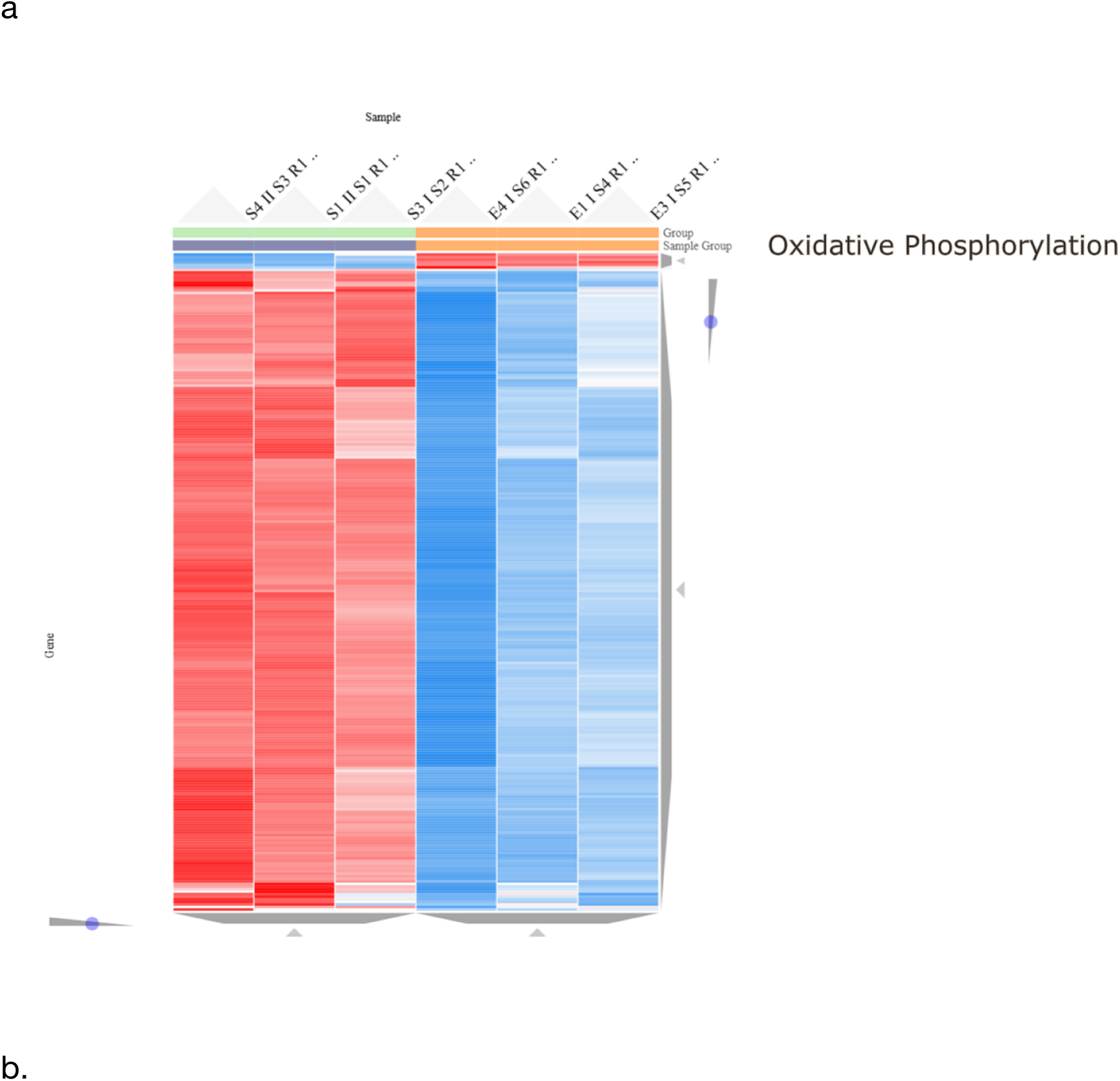
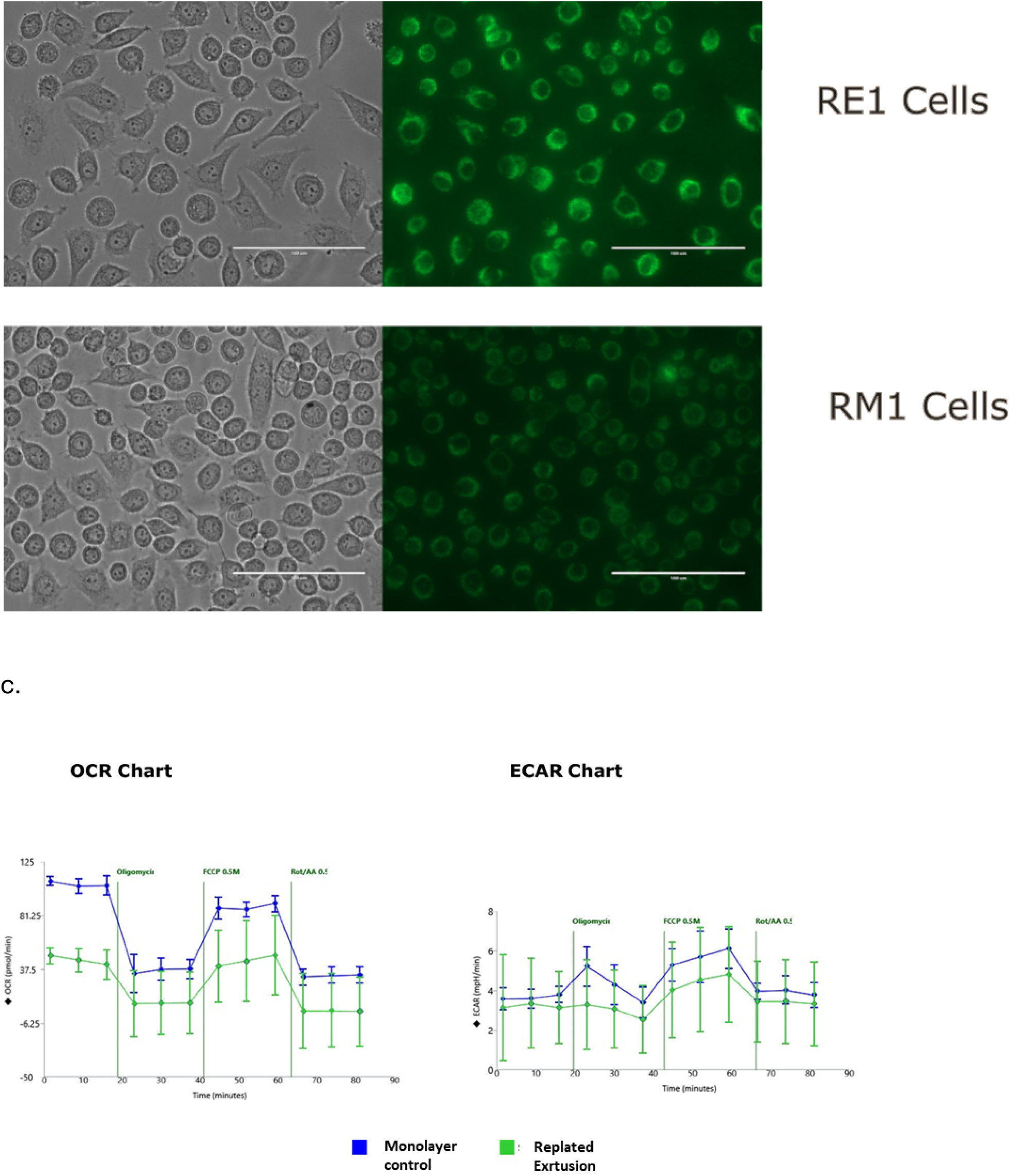
Extruded cells exhibit signs of oxidative stress response with elevated mitochondrial biogenesis and altered cellular respiration. a, Heatmap showing differential gene expression patterns related to oxidative phosphorylation across multiple samples. Red indicates upregulated genes and blue indicates downregulated genes relative to control. The samples (columns) are grouped into two distinct clusters showing opposing expression patterns. S4IIS3R1, S1IIS1R1, S3IS2R1 represent control monolayer cells and E4IS6R1, E1IS4R1, E3IS5R1 represent extruded cells. b, Comparative analysis of mitochondria in RE1 and control RM1 cells. The top row shows RE1 cells, while the bottom row shows RM1 cells. Left panels display phase-contrast images showing cellular morphology, and right panels show corresponding MitoTacker green (green) revealing the expression and localization of a mitochondria. Replated extrusion cells exhibit increased fluorescence intensity compared to the monolayer control. Scale bar = 100 μm. c, Comparison of mitochondrial function between control RM1 and RE1 cells measured by a Seahorse XF analyzer. Left panel: Oxygen Consumption Rate (OCR) demonstrating cellular respiration over time. Right panel: Extracellular Acidification Rate (ECAR) reflecting glycolytic activity. Blue lines represent RM1 cells and green lines represent RE1 cells. Sequential addition of metabolic modulators (indicated by vertical lines) was performed at the indicated time points: oligomycin (ATP synthase inhibitor, ∼20 min), FCCP (mitochondrial uncoupler, ∼40 min), and rotenone/antimycin A (Rot/AA, complex I and III inhibitors, ∼60 min). Data points represent mean values ± SEM. RM1 cells exhibit significantly higher basal respiration compared to RE1 cells, while the difference in glycolytic activity (ECAR) between the two conditions is less pronounced. This mitochondrial stress test reveals metabolic reprogramming in cells that have undergone extrusion.

**Supplementary Figure 2.**
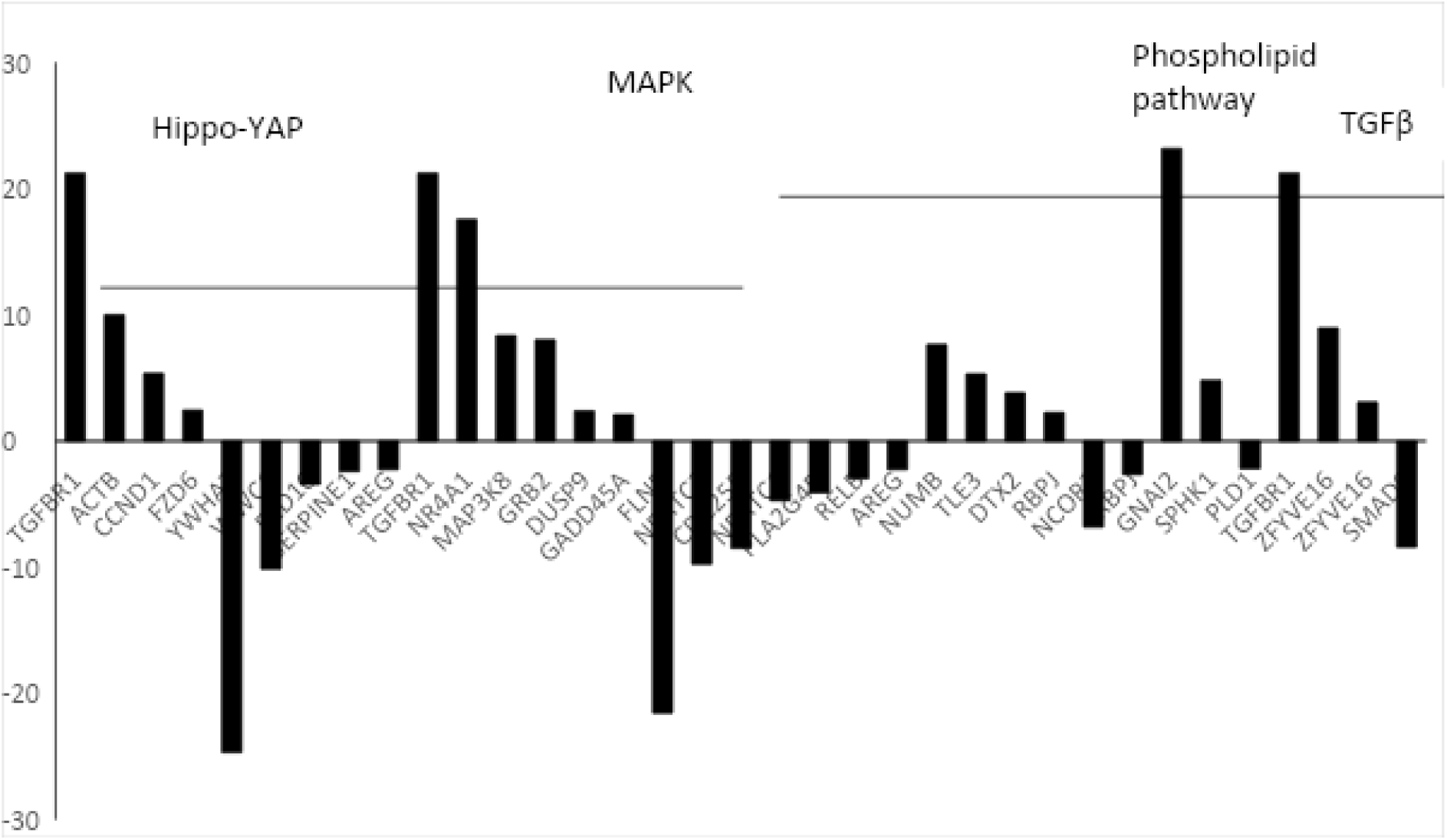
RNA sequencing of RE1 cells, compared to RM1 cells, revealed significant gene expression changes, with enrichment in phospholipid, Hippo Yap, MAPK, Notch, and TGFβ signaling pathways. a. RNA seq based gene expression analysis of RE1 vs RM1 cells showing changes related to Notch, Hippo YAP, TGFβ, MAPK and phospholipid pathways.

**Supplementary Table 1:** Relative mRNA expression of Ki67 in SIHa and HaCaT cells Ki67 fold-normalised mRNA levels (normalised to 100% confluence) in two cell lines.

| Cell line | Ki67 fold normalised mRNA (normalised to 100% confluence) |
| --- | --- |
| HaCaT | 0.38 |
| SiHa | 26.92 |

**Supplementary Table 2:**
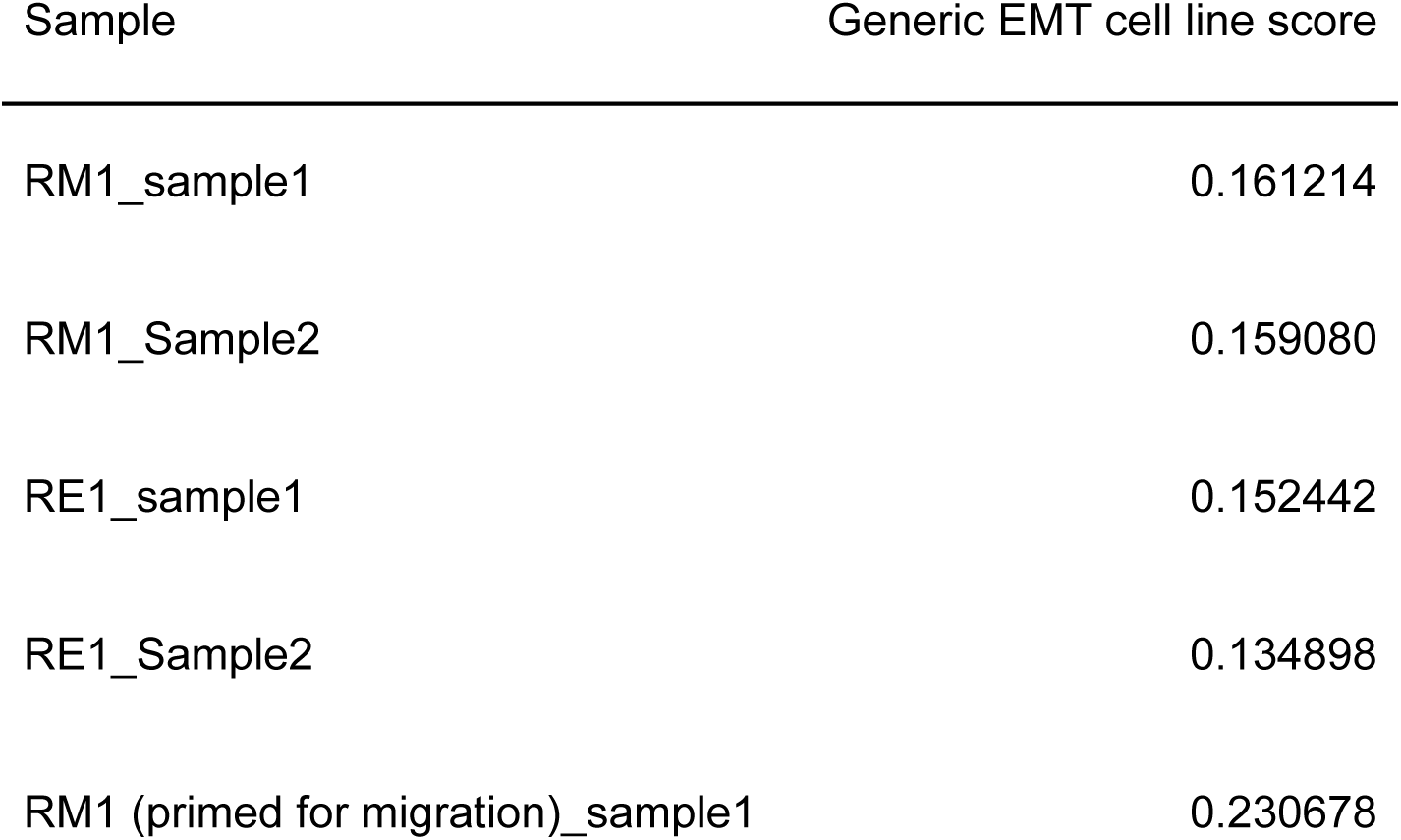
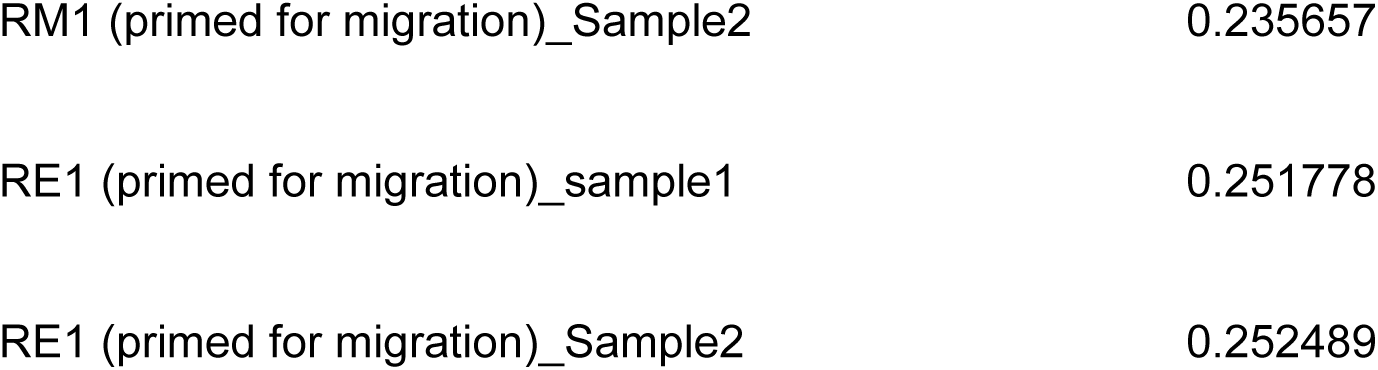
Generic EMT cell line scores across eight samples.

**Supplementary Table 3:**
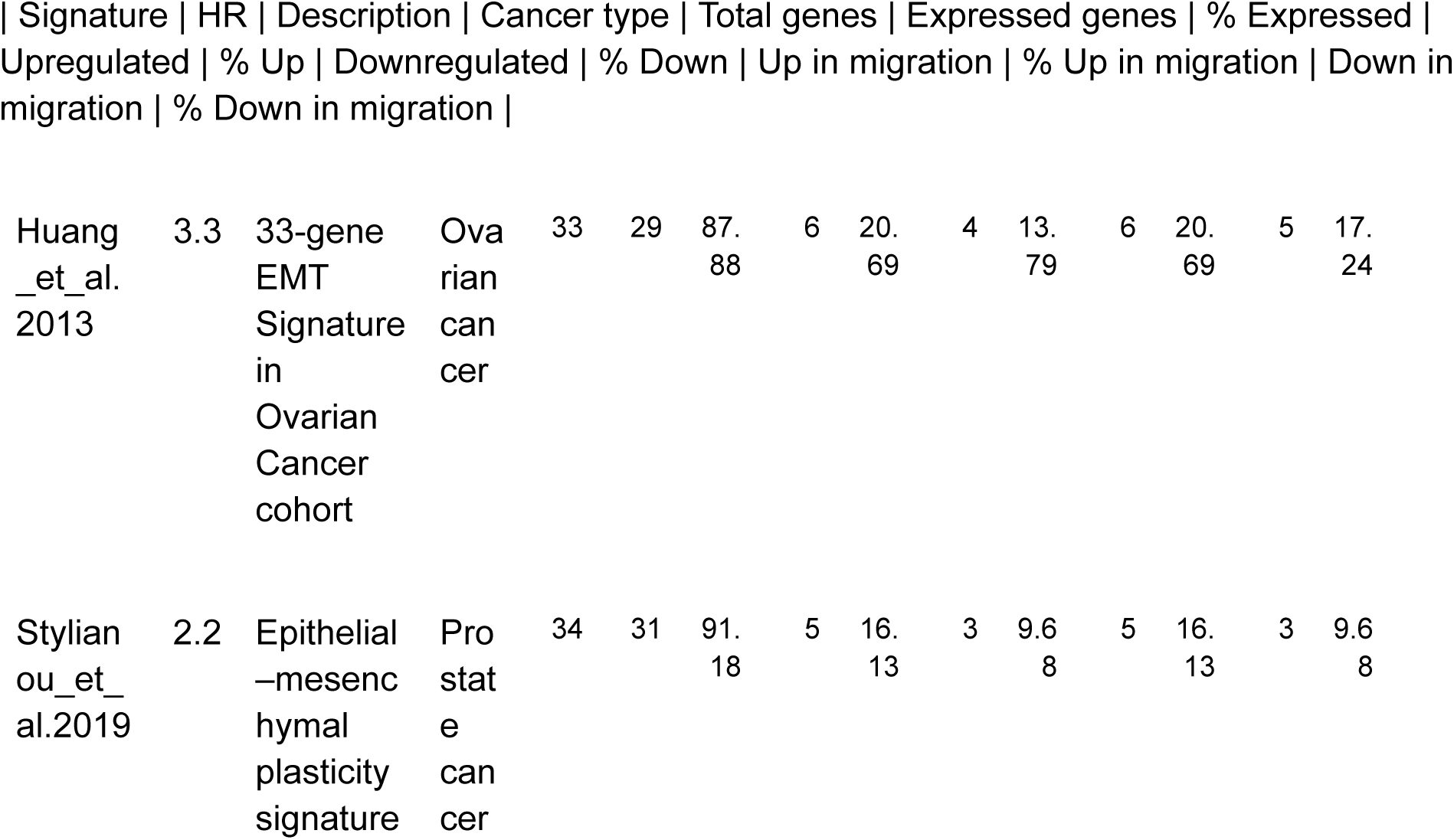
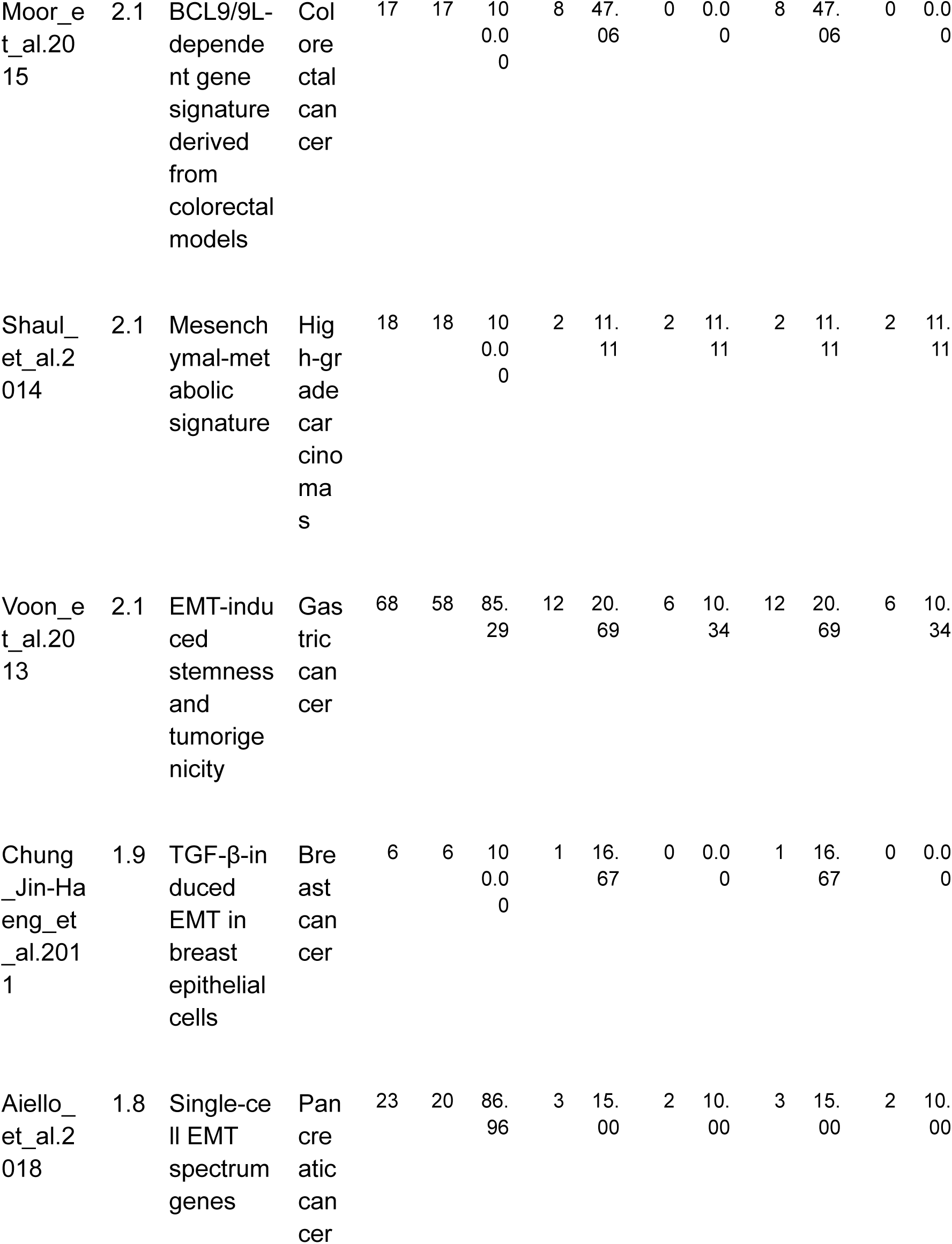
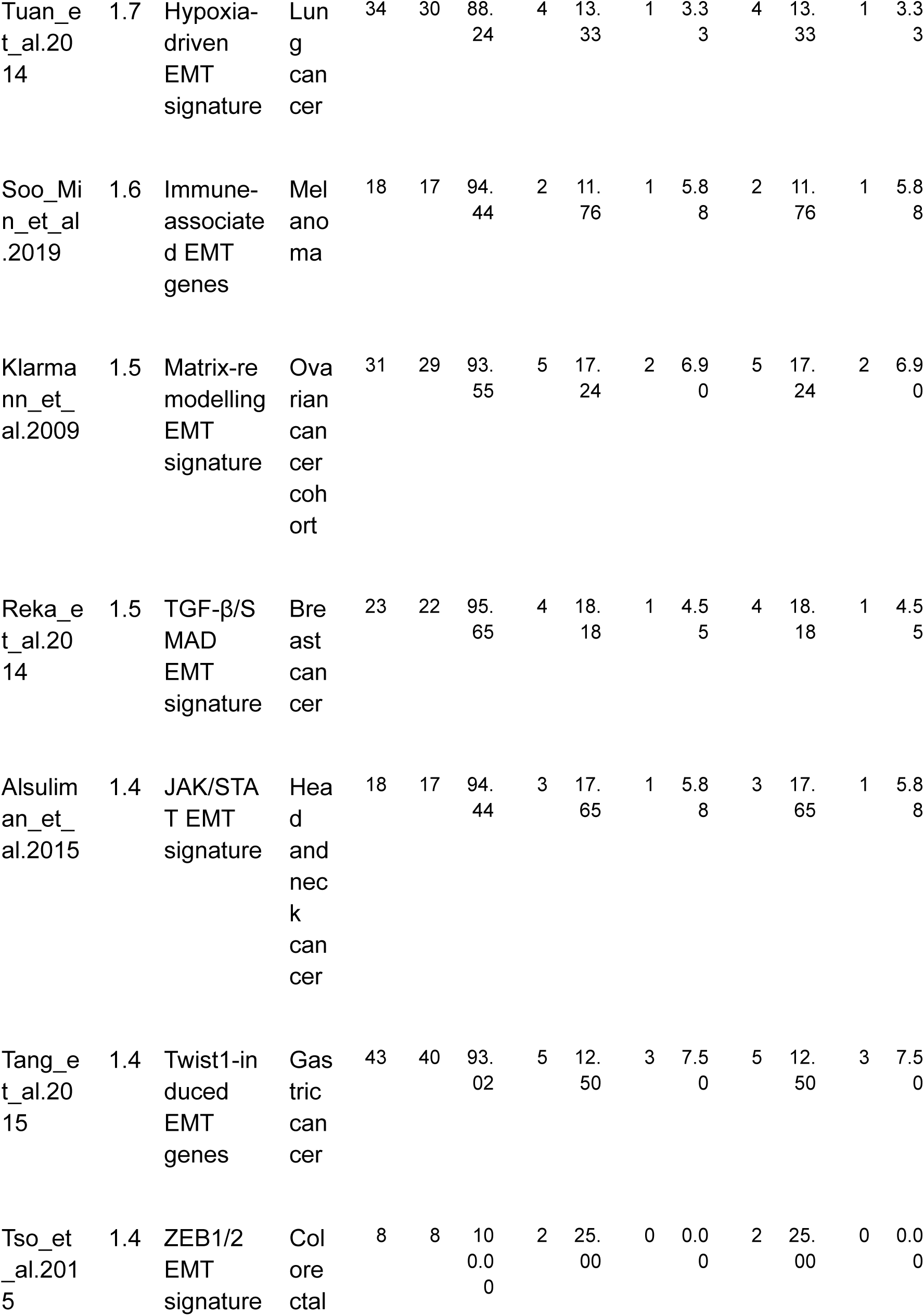
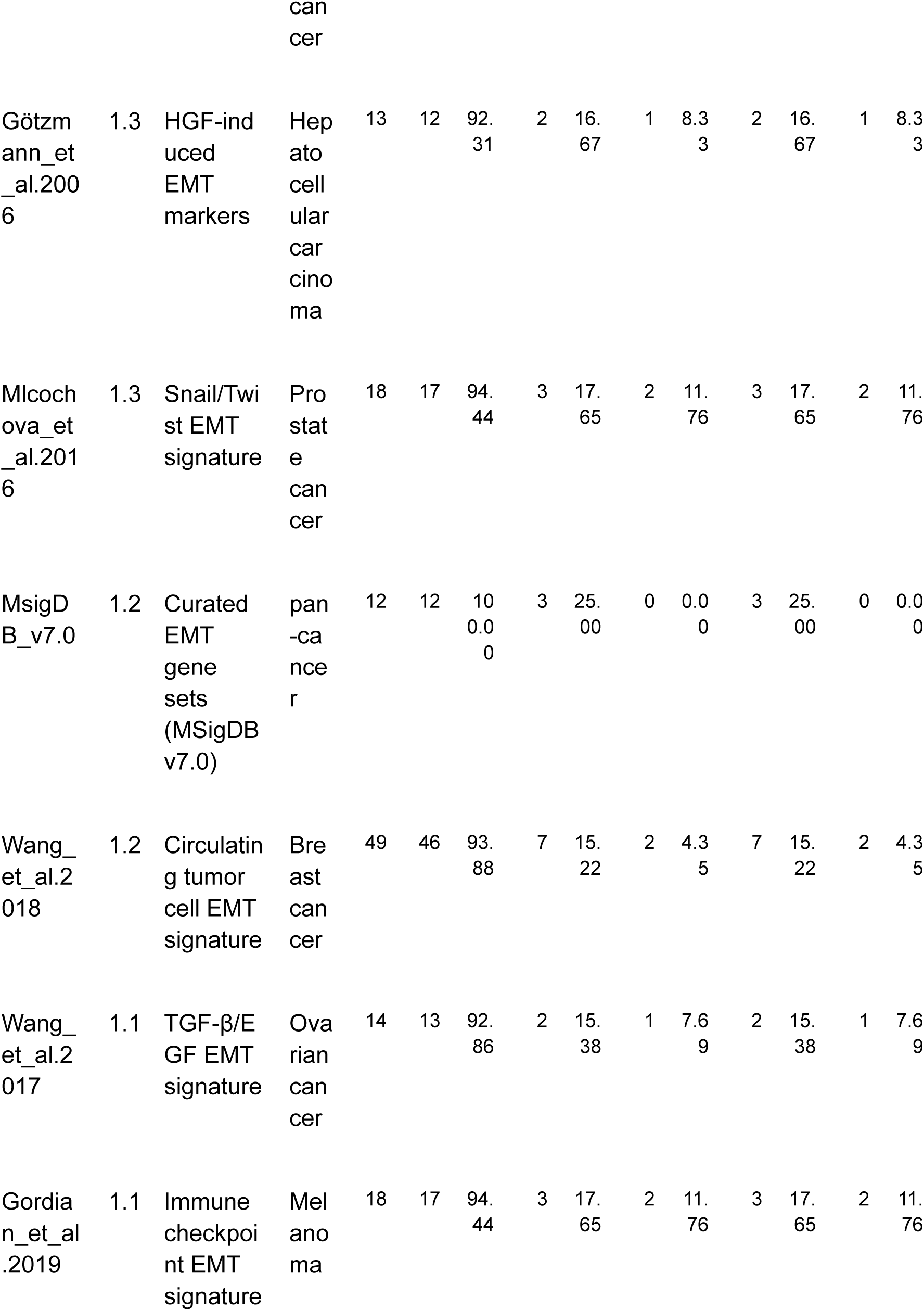
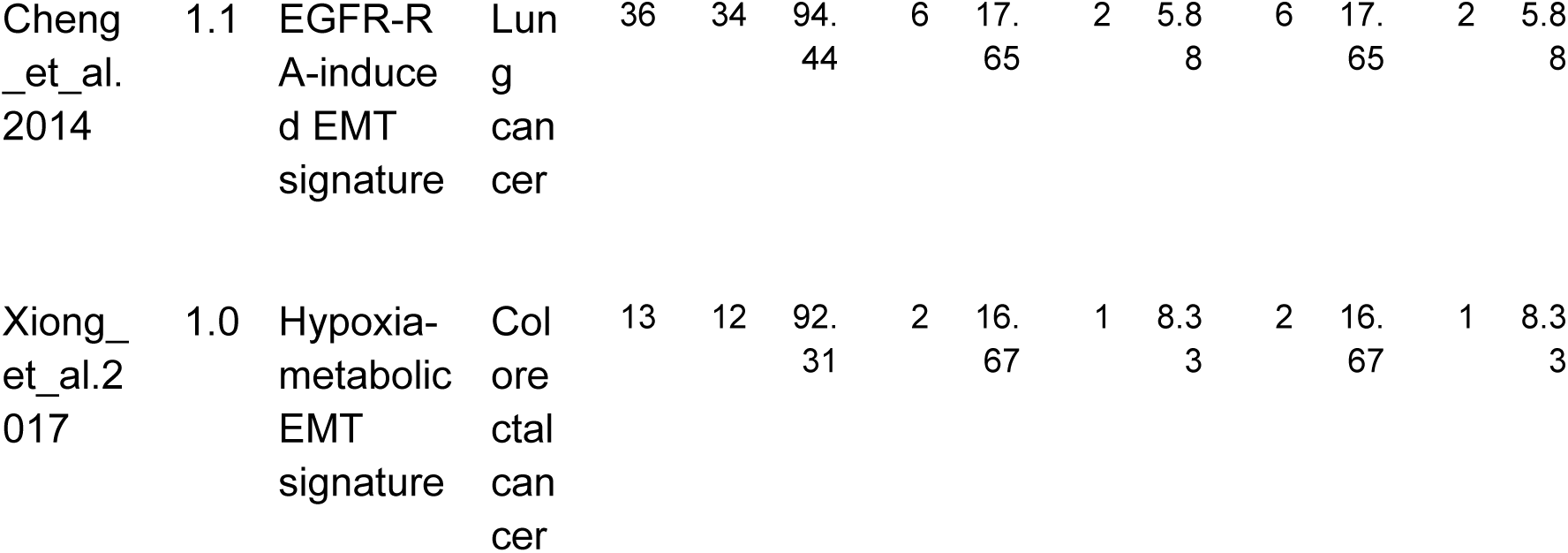
EMT Signature Hazard Ratios and Gene Expression Analysis Across Cancer Types.

